# High-level expression of STING restricts susceptibility to HBV by mediating type III IFN induction

**DOI:** 10.1101/375782

**Authors:** Hiromichi Dansako, Hirotaka Imai, Youki Ueda, Shinya Satoh, Kunitada Shimotohno, Nobuyuki Kato

## Abstract

Hepatitis B virus (HBV) is a hepatotropic DNA virus causing hepatic diseases such as chronic hepatitis, liver cirrhosis, and hepatocellular carcinoma. To study HBV, human hepatoma HepG2 cells are currently used as an HBV infectious cell culture model worldwide. HepG2 cells exhibit susceptibility to HBV by exogenously expressing sodium taurocholate cotransporting polypeptide (NTCP). We herein demonstrated that human immortalized hepatocyte NKNT-3 cells exhibited susceptibility to HBV by exogenously expressing NTCP (NKNT-3/NTCP cells). By comparing the cGAS-STING signaling pathway in several NKNT-3/NTCP cell-derived cell clones, we found that STING was highly expressed in cell clones exhibiting resistance but not susceptibility to HBV. High-level expression of STING was implicated in HBV-triggered induction of type III IFN and a pro-inflammatory cytokine, IL-6. In contrast, RNAi-mediated knockdown of STING inhibited type III IFN induction and restored the levels of HBV total transcript in an HBV-infected cell clone exhibiting resistance to HBV. These results suggest that STING regulates susceptibility to HBV by its expression levels.

STING may thus be a novel target for anti-HBV strategies.

## Introduction

Hepatitis B virus (HBV) is a hepatotropic virus classified into the *Hepadnaviridae* family. HBV infection causes chronic hepatitis, liver cirrhosis, and finally hepatocellular carcinoma (HCC) [1, 2]. The progression of hepatic diseases is tightly associated with the HBV-triggered host innate immune response and inflammatory response. To prevent the progression of hepatic diseases, it is important to suppress the HBV-triggered host innate immune response and inflammatory response.

The cytoplasmic DNA sensor, cyclic GMP-AMP synthetase (cGAS), is known to recognize viral DNA and other non-self exogenous DNAs as pathogen-associated molecular patterns (PAMPs) [3, 4]. After the recognition of non-self exogenous DNA, cGAS produces cyclic GMP-AMP (cGAMP) and then uses cGAMP to activate a stimulator of interferon genes (STING). STING mediates activation of the transcription factor interferon regulatory factor 3 (IRF-3) and subsequently the induction of interferon (IFN)-β (type I IFN) [5], IFN-λ1, λ2, and λ3 (type III IFN) [6]. Both type I and type III IFNs stimulate the induction of numerous IFN-stimulated genes (ISGs) such as ISG15 and ISG56 through the JAK-STAT signaling pathway [7]. On the other hand, STING also mediates the induction of pro-inflammatory cytokines such as IL-6 and IL-8 through the NF-κB signaling pathway [8, 9]. As described here, both cGAS and STING are required for the innate immune response and inflammatory response. We previously reported that cGAS recognized HBV DNA and subsequently triggered an innate immune response in human hepatoma Li23 cells [10]. However, in that study, we could not examine the HBV-triggered inflammatory response, since Li23 cells were a human hepatoma cell line. To study HBV-triggered inflammatory responses, it will be necessary to establish an HBV infectious cell culture model from normal human hepatic cells rather than human hepatoma cells.

Sodium taurocholate cotransporting polypeptide (NTCP) is a functional receptor for HBV [11]. Human hepatoma HepG2 cells exhibit susceptibility to HBV by exogenously expressing NTCP [11]. HepG2/NTCP cells (HepG2 cells stably expressing exogenous NTCP) are currently used as an HBV infectious cell culture model for the study of HBV worldwide. However, we previously reported that HepG2 cells exhibited defective expression of endogenous cGAS [10]. This result suggests that HepG2/NTCP cells cannot be used for the study of endogenous cGAS-triggered innate immune response and inflammatory response. Our previous study also showed that cGAS was expressed in immortalized human hepatocyte NKNT-3 cells [10]. In the present study, we established NKNT-3 cells exhibiting susceptibility to HBV by the exogenous expression of NTCP. In addition, we obtained several NKNT-3/NTCP-derived cell clones exhibiting susceptibility or resistance to HBV. Interestingly, STING was highly expressed in a cell clone exhibiting resistance to HBV. Here, we show that STING is an important host factor that regulates susceptibility to HBV by its expression levels. We also show that NKNT-3/NTCP cells are a novel HBV infectious cell culture model for the study of HBV-triggered innate immune responses and inflammatory responses.

## Results

### The immortalized human hepatocyte NKNT-3 cells exhibited susceptibility to HBV via their expression of exogenous NTCP

Since HepG2 cells were a human hepatoma cell line and exhibited defective expression of endogenous cGAS [10], we tried to establish HBV infectious cell culture model from immortalized human hepatocyte NKNT-3 cells, which has been exhibited a non-neoplastic phenotype [12] and the endogenous expression of cGAS [10]. HepG2 cells have been reported to exhibit susceptibility to HBV through their expression of exogenous NTCP [11]. Therefore, to establish NKNT-3 cells exhibiting susceptibility to HBV, we first prepared NKNT-3 cells stably expressing exogenous NTCP-myc (designated NKNT-3/NTCP cells; Fig. 1A). The cell surface expression of NTCP was detected in both NKNT-3/NTCP cells and HepG2/NTCP cells (HepG2 cells stably expressing exogenous NTCP-myc), but not in NKNT-3/Control cells (NKNT-3 cells stably expressing the control vector) (Fig. 1B). By using two kinds of inoculum, HBV/NLuc (genotype C) [13] and HBV (the supernatant of HBV-replicating HepG2.2.15 cells, genotype D) [14], we compared the levels of susceptibility to HBV in NKNT-3/NTCP cells with that in NKNT-3/Control cells. After the infection with HBV/NLuc or HBV, both level of NLuc activity and HBV total transcript were increased in NKNT-3/NTCP cells in a time-dependent manner, but not in NKNT-3/Control cells (Figs. 1C and 1D). We next compared the level of susceptibility to HBV in NKNT-3/NTCP cells with that in HepG2/NTCP cells. The levels of NLuc activity, HBV total transcript, and pgRNA in HBV/NLuc- or HBV-infected NKNT-3/NTCP cells were almost ten times lower than those in HBV/NLuc- or HBV-infected HepG2/NTCP cells (Figs. 1E and 1F). We further examined whether or not the exogenous NTCP was functional in NKNT-3/NTCP cells. Cyclosporin A (CsA) was previously reported to inhibit HBV entry by targeting NTCP [15]. When administered before and during HBV inoculation, CsA inhibited the levels of HBV total transcript in HBV-infected NKNT-3/NTCP cells as well as in HBV-infected HepG2/NTCP cells (Fig. 1G). These results suggest that NKNT-3 cells exhibit susceptibility to HBV by exogenously expressing functional NTCP.

**Figure 1.**
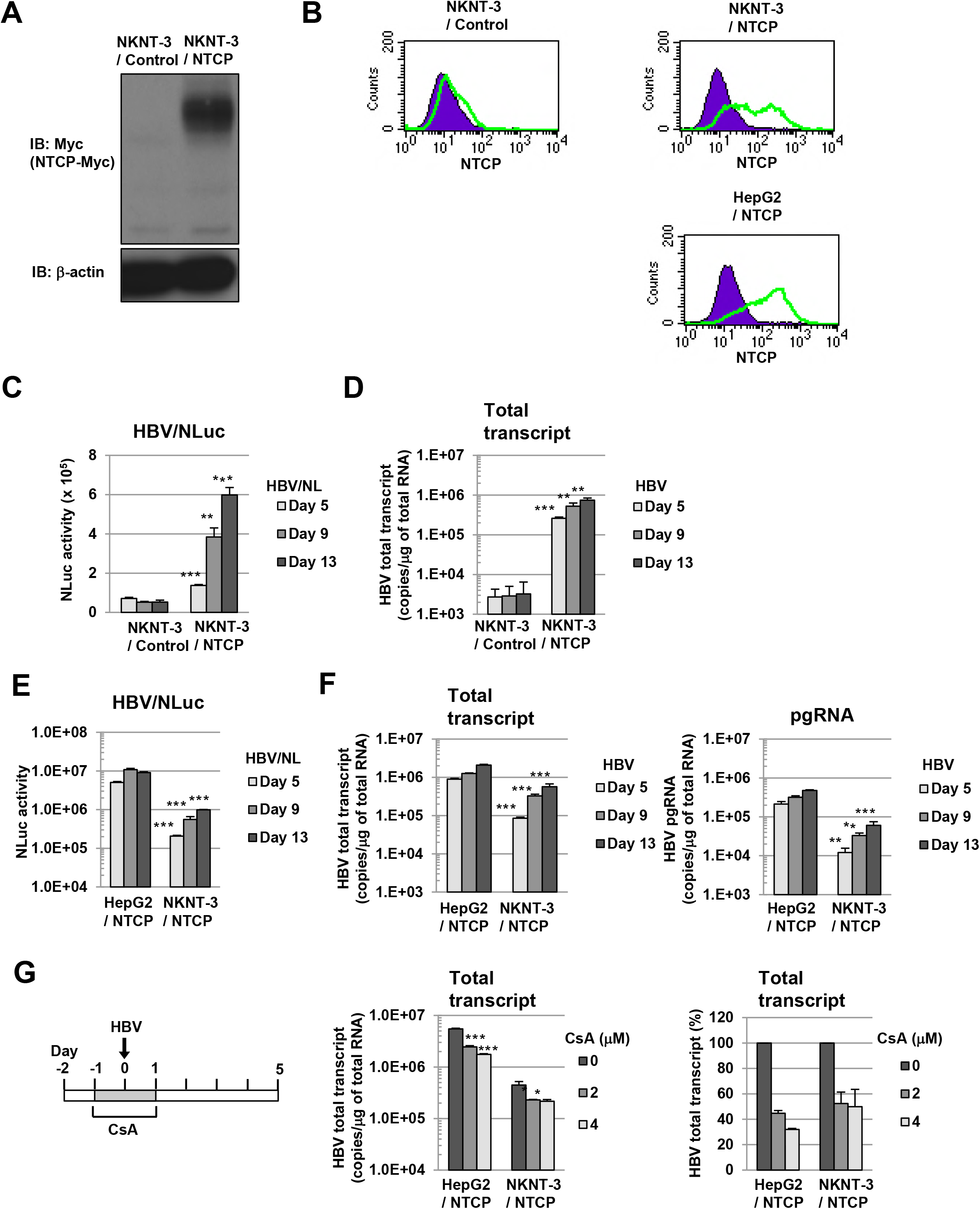
The immortalized human hepatocyte cell line NKNT-3 exhibited susceptibility to HBV by expressing exogenous NTCP. A Western blot analysis of exogenous NTCP in NKNT-3/NTCP cells. Anti-Myc antibody was used for the detection of NTCP-Myc in NKNT-3/NTCP cells. β-actin was included as a loading control. B Flow cytometric analysis of the cell surface NTCP in NKNT-3/Control cells, NKNT-3/NTCP cells, or HepG2/NTCP cells. Signals of the cell surface NTCP are shown in green. An isotype control was used as a negative control (violet area). C Comparison of NLuc activity after HBV/NL inoculation between NKNT-3/Control cells and NKNT-3/NTCP cells. Intracellular NLuc activity was measured at 5, 9, and 13 days after HBV/NL inoculation. \*\**P* < 0.01, \*\*\**P* < 0.001 versus HBV/NL-infected NKNT-3/Cont cells. D Quantitative RT-PCR analysis of the amount of HBV total transcript in HBV-infected NKNT-3/Control cell or NKNT-3/NTCP cells. The supernatant of HepG2.2.15 cells was used as an HBV inoculum. The amounts of HBV total transcript were measured at 5, 9, and 13 days after HBV inoculation. \*\**P* < 0.01, \*\*\**P* < 0.001 versus HBV-infected NKNT-3/Cont cells. E, F Comparison of the susceptibility to HBV between HepG2/NTCP cells and NKNT-3/NTCP cells. Intracellular NLuc activity was measured after HBV/NL inoculation. The amounts of HBV total transcript and the pgRNA were measured after HBV inoculation by quantitative RT-PCR analysis. \*\**P* < 0.01, \*\*\**P* < 0.001 versus HBV/NL- or HBV-infected HepG2/NTCP cells, respectively. G Functional analysis of NTCP in NKNT-3/NTCP cells using CsA as an HBV-entry inhibitor. CsA was administered before and during HBV inoculation. \**P* < 0.05, \*\*\**P* < 0.001 versus 0 μM of CsA-administered HBV-infected cells.

### The level of susceptibility to HBV in NKNT-3/NTCP #28.3.8 cells approximated that in HepG2/NTCP cells

Since susceptibility to HBV in NKNT-3/NTCP cells was lower than that in HepG2/NTCP cells (Figs. 1E and 1F), we next tried to select a subcloned cell line exhibiting higher susceptibility to HBV than NKNT-3/NTCP cells (Fig. 2A). During three-round serial limited dilution, we obtained three distinct cell clones (#28, #28.3, and #28.3.8 cells, respectively; Fig. 2A) that met this criterion (Fig. 2B). Exogenous NTCP was expressed on the cell surface in all three clones (Fig. 2C). Among them, the NKNT-3/NTCP #28.3.8 cells exhibited the highest levels of HBV total transcript after HBV infection (Fig. 2D). Therefore, we next compared the levels of susceptibility to HBV in NKNT-3/NTCP #28.3.8 cells with those in HepG2/NTCP cells. Upon the infection with HBV/NLuc or HBV, both levels of NLuc activity (Fig. 2E) and HBV total transcript (Fig. 2F) in NKNT-3/NTCP #28.3.8 cells approximated those in HepG2/NTCP cells. Consistent with these results, Northern blot analysis also showed that the levels of HBV pregenomic RNA (pgRNA) and 2.1/2.3 kb RNA in NKNT-3/NTCP #28.3.8 cells were roughly the same as those in HepG2/NTCP cells after HBV infection (Fig. 2G). These results suggest that NKNT-3/NTCP #28.3.8 cells are useful as an HBV infectious cell culture model in the manner of HepG2/NTCP cells.

**Figure 2.**
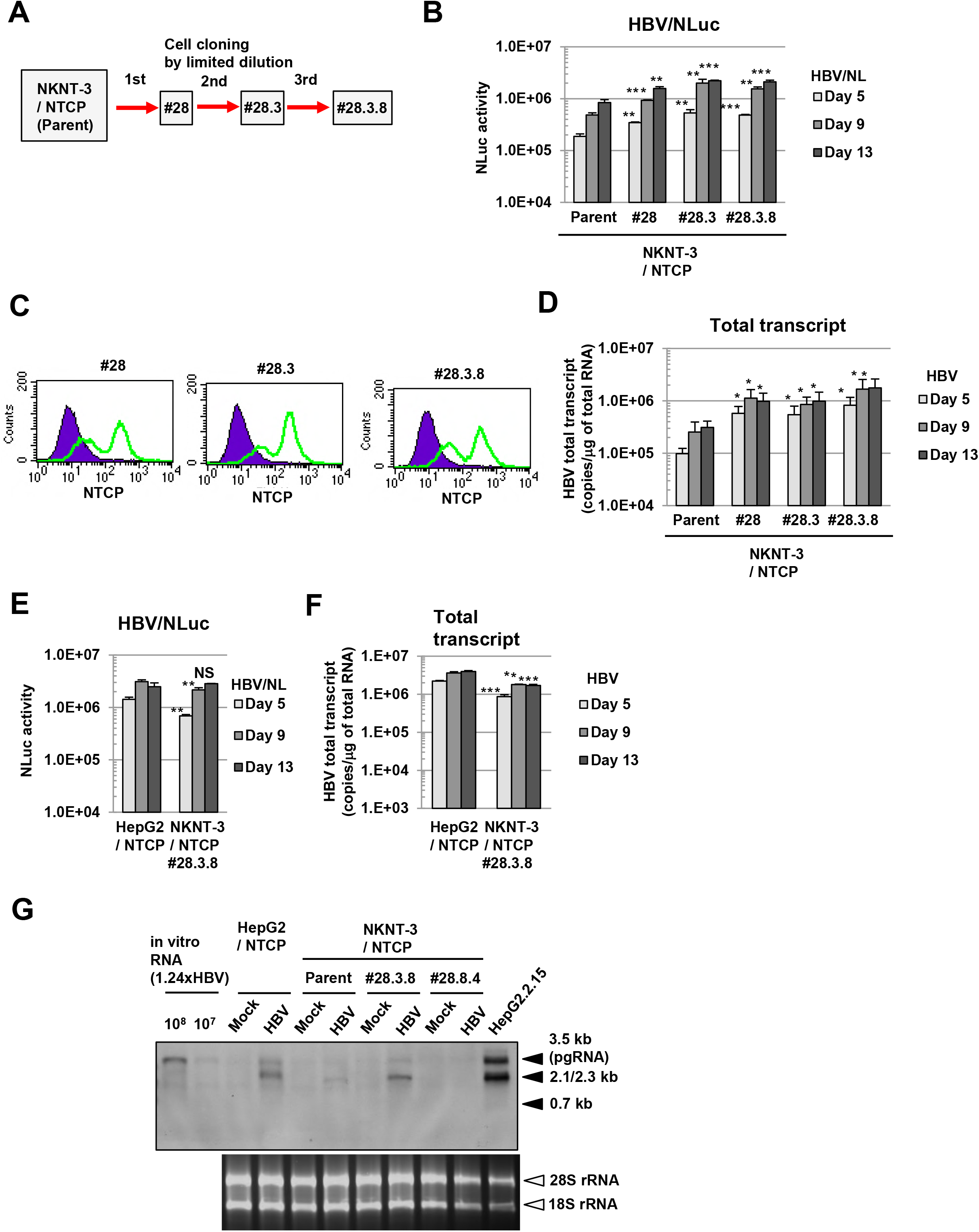
The level of susceptibility to HBV in NKNT-3/NTCP #28.3.8 cells approximated that in HepG2/NTCP cells. A Outline of cell cloning by the limited dilution method. NKNT-3/NTCP #28.3.8 cells were selected by three-round limited dilution. Red arrows with solid lines show the selection of a cell clone exhibiting higher susceptibility to HBV B Comparison of susceptibility to HBV among parent NKNT-3/NTCP cells and their derived cell clones by using HBV/NL assay. \*\**P* < 0.01, \*\*\**P* < 0.001 versus HBV/NL-infected parent NKNT-3/NTCP cells. C Flow cytometric analysis of the cell surface NTCP in their derived cell clones. Signals of the cell surface NTCP are shown in green. An isotype control was used as a negative control (violet area). D Comparison of the amounts of HBV total transcript after HBV infection among parent NKNT-3/NTCP cells and their derived cell clones. The amount of HBV total transcript was measured after HBV infection by quantitative RT-PCR analysis. \**P* < 0.05 versus HBV-infected parent NKNT-3/NTCP cells. E, F Comparison of susceptibility to HBV between HepG2/NTCP cells and NKNT-3/NTCP #28.3.8 cells. Intracellular NLuc activity or the amounts of HBV total transcript were measured as described in Figs. 1E and 1F. NS; not significant, \*\**P* < 0. 01, \*\*\**P* < 0.001 versus HBV/NL- or HBV-infected HepG2/NTCP cells, respectively. G Comparison of susceptibility to HBV between HepG2/NTCP cells and NKNT-3/NTCP #28.3.8 cells by Northern blot analysis. Total RNA was isolated from HBV-infected cells at 13 days after HBV inoculation. 28S rRNA and 18S rRNA were included as a loading control. NKNT-3/NTCP #28.8.4 is another clone, which has been estimated to exhibit susceptibility to HBV by HBV/NL assay (data not shown).

### HBV triggered the induction of type III IFNs in NKNT-3/NTCP #28.3.25.13 cells exhibiting resistance to HBV

During the three-round limited dilution, we obtained NKNT-3/NTCP #28.3.8 cells that exhibited higher susceptibility to HBV than the parent NKNT-3/NTCP cells (Figs. 2B and 2D). On the other hand, during the additional limited dilution (Fig. 3A), we unexpectedly obtained a cell clone (#28.3.25.13) exhibiting greater resistance to HBV compared with NKNT-3/NTCP #28.3.8 cells (Fig. 3B). We conjectured that the innate immune response might be induced in cell clones exhibiting resistance to HBV. To examine this possibility, we first compared the HBV-triggered innate immune responses among cell clones exhibiting susceptibility or resistance to HBV. At 5 days after HBV infection, ISG56 was strongly induced in NKNT-3/NTCP #28.3.25.13 cells, but not in NKNT-3/NTCP #28.3.8 cells (Fig. 3C). Since HBV-triggered ISG56 induction in NKNT-3/NTCP #28.3.25.13 cells was higher than that in #28.3. 30.20.3 cells (another cell clone exhibiting resistance to HBV, Fig. 3B), we mainly focused the innate immune response to HBV in NKNT-3/NTCP #28.3.25.13 cells. We first compared the time course of *ISG56* mRNA induction after HBV infection between NKNT-3/NTCP #28.3.8 and #28.3.25.13 cells (Fig. 3D). At 5 or 9 days after HBV infection, *ISG56* mRNA was strongly induced in NKNT-3/NTCP #28.3.25.13 cells, but not in #28.3.8 cells (Fig. 3D).

**Figure 3.**
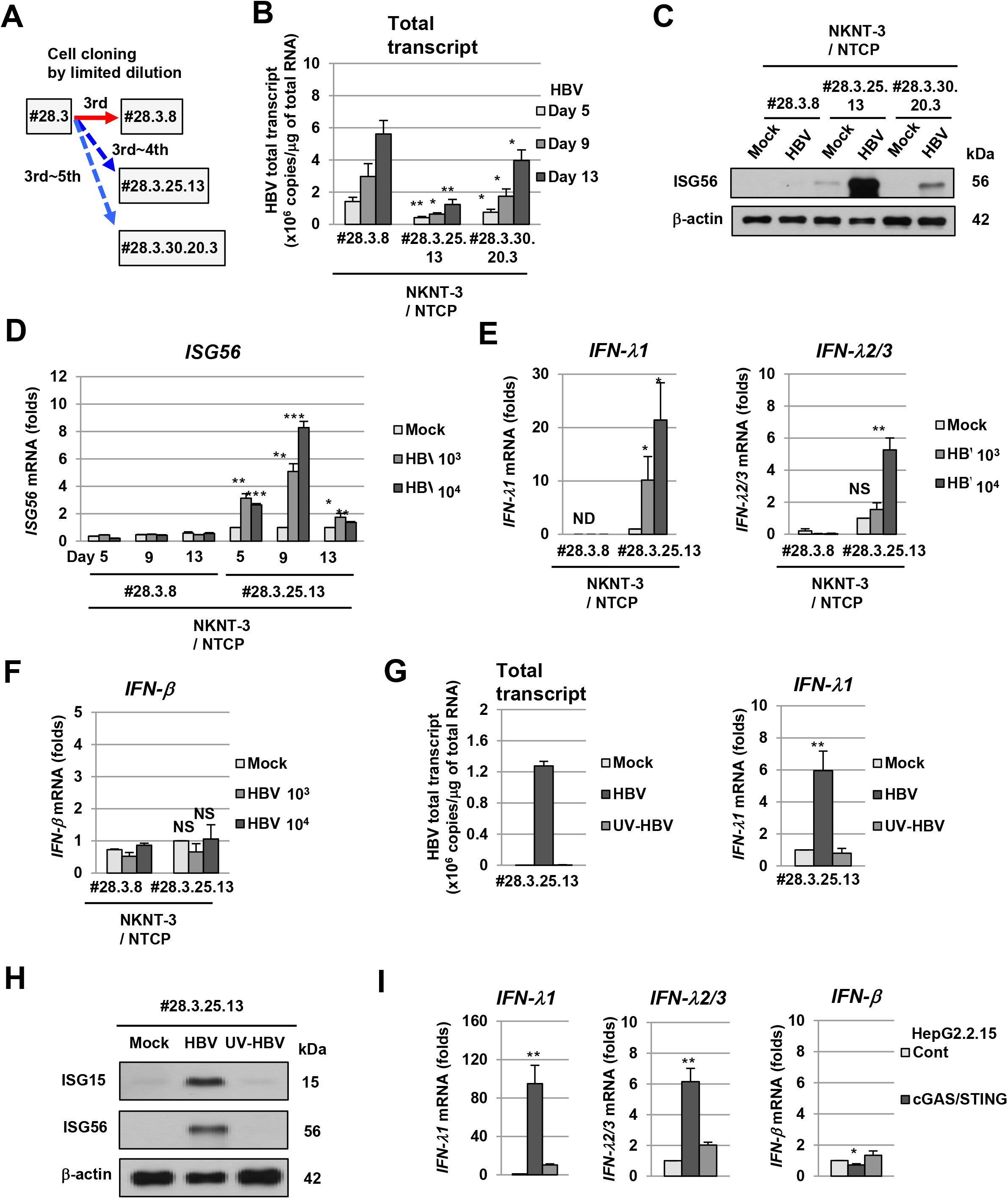
HBV induced type III IFN in NKNT-3/NTCP #28.3.25.13 cells exhibiting resistance to HBV. A Outline of cell cloning by the limited dilution method. NKNT-3/NTCP #28.3.25.13 and #28.3.30.20.3 cells were selected by their distinct serial limited dilution, respectively. Blue arrows with dashed lines show the selection of a cell clone exhibiting resistance to HBV. B Quantitative RT-PCR analysis of the amounts of HBV total transcript in HBV-infected NKNT-3/NTCP #28.3.8, #28.3.25.13, or #28.3.30.20.3 cells. \**P* < 0.05, \*\**P* < 0.01 versus HBV-infected NKNT-3N #28.3.8 cells. C Western blot analysis of ISG56 in HBV-infected NKNT-3/NTCP #28.3.8, #28.3.25.13, or #28.3.30.20.3 cells. Cell lysates were prepared from mock- or HBV-infected cells at 5 days after HBV inoculation. D Quantitative RT-PCR analysis of *ISG56* mRNA in HBV-infected NKNT-3/NTCP #28.3.8 or #28.3.25.13 cells. Cells were infected with HBV at 10^3^ or 10^4^ HBV genome equivalents per cell, respectively. Each mRNA level was calculated relative to the level in mock-infected NKNT-3/NTCP #28.3.25.13 cells, which was set at 1. \**P* < 0.05, \*\**P* < 0.01, \*\*\**P* < 0.001 versus mock-infected NKNT-3N #28.3.25.13 cells. E Quantitative RT-PCR analysis of *IFN-λ1* and *IFN-λ2/3* mRNA in HBV-infected NKNT-3/NTCP #28.3.8 or #28.3.25.13 cells. Cells were infected with HBV at 10^3^ or 10^4^ HBV genome equivalents per cell, respectively. Each mRNA level was calculated as described in Fig. 3D. ND, not detected. NS; not significant, \**P* < 0.05, \*\**P* < 0.01 versus mock-infected NKNT-3/NTCP #28.3.25.13 cells. F Quantitative RT-PCR analysis of *IFN-β* mRNA in HBV-infected NKNT-3/NTCP #28.3.8 or #28.3.25.13 cells. Cells were infected with HBV at 10^3^ or 10^4^ HBV genome equivalents per cell, respectively. Each mRNA level was calculated as described in Fig. 3D. NS; not significant versus mock-infected NKNT-3/NTCP #28.3.25.13 cells. G (left panel) Quantitative RT-PCR analysis of the amounts of HBV total transcript in mock-, HBV-, or UV-HBV-infected NKNT-3/NTCP #28.3.25.13 cells. (right panels) Quantitative RT-PCR analysis of *IFN-λ1* mRNA in mock-, HBV-, or UV-HBV-infected NKNT-3/NTCP #28.3.25.13 cells. Each mRNA level was calculated as described in Fig. 3D. \*\**P* < 0.01 versus mock- or UV-HBV-infected NKNT-3/NTCP #28.3.25.13 cells, respectively. H Western blot analysis of ISG15 and ISG56 in mock-, HBV-, or UV-HBV-infected NKNT-3/NTCP #28.3.25.13 cells. The cell lysate was prepared as described in Fig. 3C. I Quantitative RT-PCR analysis of *IFN-λ1, IFN-λ2/3*, and *IFN-β* mRNA in HepG2.2.15 cGAS/STING cells. Each mRNA level was calculated relative to the level in HepG2.2.15 Cont cells, which was set at 1. \**P* < 0.05, \*\**P* < 0.01 versus HepG2.2.15 Cont cells or HepG2.2.15 cGAS GSAA/STING cells, respectively.

These results suggest that HBV infection induces the innate immune response in cell clone exhibiting resistance but not susceptibility to HBV. We next examined whether type I and/or type III IFN was required for *ISG56* mRNA induction after HBV infection in NKNT-3/NTCP #28.3.25.13 cells. Interestingly, at 9 days after HBV infection, *IFN-λ1* and *IFN-λ2/3* (type III IFN) mRNA, but not *IFN-β* (type I IFN) mRNA, were induced in NKNT-3/NTCP #28.3.25.13 cells (Figs. 3E and 3F). In addition, *IFN-λ1* mRNA (Fig. 3G), ISG15 (Fig. 3H), and ISG56 (Fig. 3H) were induced at 9 days after HBV infection, but not mock or ultraviolet-inactivated HBV (UV-HBV) infection, in NKNT-3/NTCP #28.3.25.13 cells. Consistent with these results, HBV induced *IFN-λ1* and *IFN-λ2/3*, but not *IFN-β* mRNA, in HBV-replicating HepG2.2.15 cGAS/STING cells stably expressing both exogenous cGAS and STING [10] (Fig. 3I). In addition, the induction levels of *IFN-λ1* and *IFN-λ2/3* mRNA in HepG2.2.15 cGAS/STING cells were higher than those in HepG2.2.15 cGAS GSAA/STING cells stably expressing both exogenous cGAS GSAA (the inactive mutant of cGAS) and STING [10]. These results suggest that HBV induces type III IFN through the cGAS/STING signaling pathway in NKNT-3/NTCP #28.3.25.13 cells, but not in #28.3.8 cells. These results also suggest that the expression levels of cGAS/STING signaling pathway-associated host factor(s) are different between NKNT-3/NTCP #28.3.8 cells and #28.3.25.13 cells.

### High-level expression of STING was implicated in HBV-triggered type III IFN induction in NKNT-3/NTCP #28.3.25.13 cells exhibiting resistance to HBV

Since our results suggested that the expression levels of cGAS/STING signaling pathway-associated host factor(s) were different between NKNT-3/NTCP #28.3.8 cells and #28.3.25.13 cells, we next compared the levels of p-dGdC (the synthetic ligand for the cGAS/STING signaling pathway)-triggered type III IFN induction. We found that the p-dGdC-triggered *ISG56* and *IFN-lambdal* mRNA induction in NKNT-3/NTCP #28.3.25.13 cells was several times higher than that in NKNT-3/NTCP #28.3.8 cells (Fig. 4A). We next tried to identify the host factor(s) responsible for the higher responsiveness to p-dGdC in NKNT-3/NTCP #28.3.25.13 cells. Among cGAS/STING signaling pathway-associated host factor(s), we found that *STING* mRNA (Fig. 4B) and STING protein (Fig. 4C) were highly expressed in NKNT-3/NTCP #28.3.25.13 cells. These results suggest that the high-level expression of STING enhances p-dGdC-triggered type III IFN induction in NKNT-3/NTCP #28.3.25.13 cells compared with #28.3.8 cells. We further compared the phosphorylation levels of STING among several NKNT-3/NTCP cell-derived cell clones. STING was highly phosphorylated in p–dGdC-transfected NKNT-3/NTCP #28.3.25.13 cells, but not in #28.3.8 cells (Fig. 4D, lower-left panel). In addition, STING was also highly phosphorylated in p–dGdC-treated NKNT-3/NTCP #28.3.25 cells (the parent cells of #28.3.25.13) but not in the parent cells, or in #28 and #28.3 cells (the common parent cells of #28.3.8, #28.3.25 and #28.3.25.13, respectively). *IFN-λ1* mRNA was strongly induced in NKNT-3/NTCP cells highly phosphorylating STING such as NKNT-3/NTCP #28.3.25 and #28.3.25.13 cells (Fig. 4D, upper-left panel). Consistent with these results, the knockdown of STING reduced *IFN-λ1* mRNA induction in p-dGdC-transfected NKNT-3/NTCP #28.3.25.13 cells (Fig. 4D, upper-right panel). These results suggest that STING regulate p-dGdC-triggered type III IFN induction by its expression level in NKNT-3/NTCP cells.

**Figure 4.**
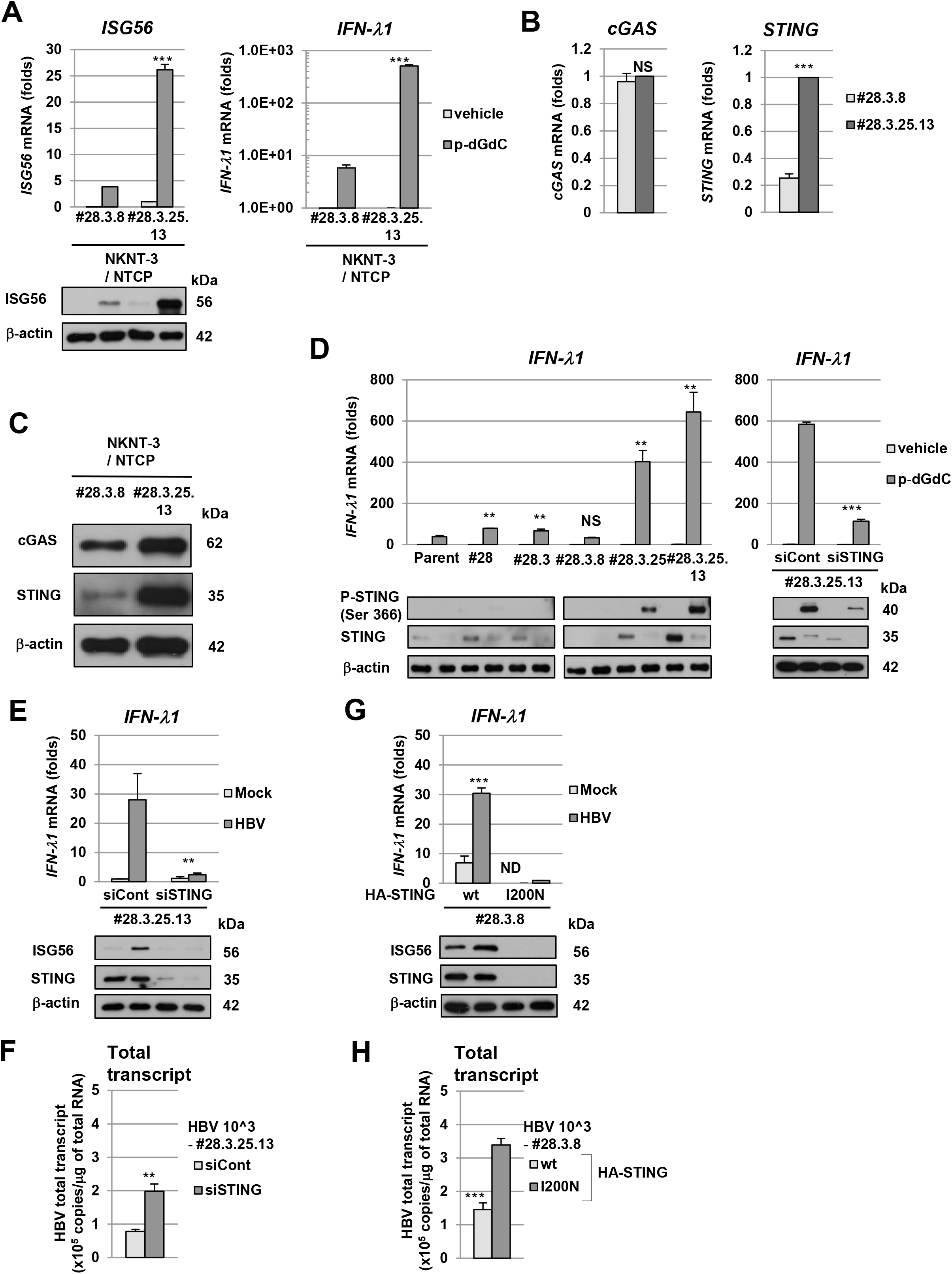
High-level expression of STING was implicated in HBV-triggered type III IFN induction in NKNT-3/NTCP #28.3.25.13 cells. A (upper panel) Quantitative RT-PCR analysis of *ISG56* and *IFN-λ1* mRNA in p-dGdC-transefected NKNT-3/NTCP #28.3.8 or #28.3.25.13 cells. Each mRNA level was calculated relative to the level in vehicle-transfected NKNT-3/NTCP #28.3.25.13 cells, which was set at 1. \*\*\**P* <sg 0.001 versus p-dGdC-transfected NKNT-3/NTCP #28.3.8 cells. (lower panel) Western blot analysis of ISG56 in p-dGdC-transefected NKNT-3/NTCP #28.3.8 or #28.3.25.13 cells. The cell lysate was prepared as described in Fig. 3C. B Quantitative RT-PCR analysis of *cGAS* and *STING* mRNA in NKNT-3/NTCP #28.3.8 or #28.3.25.13 cells. Each mRNA level was calculated relative to the level in NKNT-3/NTCP #28.3.25.13 cells, which was set at 1. NS; not significant, \*\*\**P* < 0.001 versus NKNT-3/NTCP #28.3.8 cells. C Western blot analysis of cGAS and STING in NKNT-3/NTCP #28.3.8 or #28.3.25.13 cells. D (upper-left panel) Quantitative RT-PCR analysis of *IFN-λ1* mRNA in the parent NKNT-3/NTCP cells and in the several cell clones derived from them after transfection with p-dGdC. Each mRNA level was calculated relative to the level in NKNT-3/NTCP #28.3.25.13 cells transfected with vehicle, which was set at 1. NS; not significant, \*\**P* < 0.01 versus p-dGdC-transfected parent NKNT-3/NTCP cells. (lower-left panel) Western blot analysis of phosphorylated STING at Ser366 in the original NKNT-3/NTCP cells and in the several cell clones derived from them after transfection with p-dGdC. The cell lysate was prepared as described in Fig. 3C. (upper-right panel) Quantitative RT-PCR analysis of *IFN-λ1* mRNA in NKNT-3/NTCP #28.3.25.13 cells transfected with STING-specific (designated NKNT-3/NTCP #28.3.25.13 siSTING) or control (designated NKNT-3/NTCP #28.3.25.13 siCont) siRNA followed by p-dGdC. Each mRNA level was calculated relative to the level in vehicle-transfected NKNT-3/NTCP #28.3.25.13 siCont cells, which was set at 1. \*\*\**P* < 0.001 versus p-dGdC-transfected NKNT-3/NTCP #28.3.25.13 siCont cells. (lower-right panel) Western blot analysis of phosphorylated STING at Ser366 in NKNT-3/NTCP #28.3.25.13 siSTING cells after transfection with p-dGdC. The cell lysate was prepared as described in Fig. 3C. E (upper panel) Quantitative RT-PCR analysis of *IFN-λ1* mRNA in mock- or HBV-infected NKNT-3/NTCP #28.3.25.13 siSTING cells or NKNT-3/NTCP #28.3.25.13 siCont cells. Each mRNA level was calculated relative to the level in mock-infected NKNT-3/NTCP #28.3.25.13 siCont cells, which was set at 1. (lower panel) Western blot analysis of ISG56 in HBV-infected NKNT-3/NTCP #28.3.25.13 siCont cells or NKNT-3/NTCP #28.3.25.13 siSTING cells. The cell lysate was prepared as described in Fig. 3C. \*\**P* < 0.01 versus HBV-infected NKNT-3/NTCP #28.3.25.13 siCont cells. F Quantitative RT-PCR analysis of the amount of HBV total transcript in HBV-infected NKNT-3/NTCP #28.3.25.13 siCont cells or NKNT-3/NTCP #28.3.25.13 siSTING cells. \*\**P* < 0.01 versus HBV-infected NKNT-3/NTCP #28.3.25.13 siCont cells. G (upper panel) Quantitative RT-PCR analysis of *IFN-λ1* mRNA in mock- or HBV-infected NKNT-3/NTCP #28.3.8 cells stably expressing exogenous STING wild type (designated NKNT-3/NTCP #28.3.8 STING wt) or STING I200N (designated NKNT-3/NTCP #28.3.8 STING I200N). Each mRNA level was calculated relative to the level in HBV-infected NKNT-3/NTCP #28.3.8 STING I200N cells, which was set at 1. ND, not detected. \*\*\**P* < 0.001 versus HBV-infected NKNT-3/NTCP #28.3.8 STING I200N cells. (lower panel) Western blot analysis of ISG56 in HBV-infected NKNT-3/NTCP #28.3.8 STING wt cells or NKNT-3/NTCP #28.3.8 STING I200N cells. The cell lysate was prepared as described in Fig. 3C. H Quantitative RT-PCR analysis of the amount of HBV total transcript in HBV-infected NKNT-3/NTCP #28.3.8 STING wt cells or NKNT-3/NTCP #28.3.8 STING I200N cells. \*\*\**P* < 0.001 versus HBV-infected NKNT-3/NTCP #28.3.8 STING I200N cells.

We next examined whether high-level expression of STING was required for HBV-triggered type III IFN induction in NKNT-3/NTCP #28.3.25.13 cells. We found that knockdown of STING decreased the induction of *IFN-λl* mRNA (Fig. 4E, upper panel) and subsequently ISG56 (Fig. 4E, lower panel) in HBV-infected NKNT-3/NTCP #28.3.25.13 cells. The knockdown of STING also increased the amounts of HBV total transcript in HBV-infected NKNT-3/NTCP #28.3.25.13 cells (Fig. 4F). On the other hand, the stable expression of exogenous STING, but not STING I200N which causes the conformational disruption [16], increased the induction of *IFN-λl* mRNA (Fig. 4G, upper panel) and subsequently ISG56 (Fig. 4G, lower panel) in HBV-infected NKNT-3/NTCP #28.3.8 cells. The stable expression of exogenous STING also decreased the amounts of HBV total transcript in HBV-infected NKNT-3/NTCP #28.3.8 cells (Fig. 4H). These results suggest that high-level expression of STING is implicated in HBV-triggered type III IFN induction in NKNT-3/NTCP #28.3.25.13 cells.

### High-level expression of STING was required for the HBV-triggered inflammatory response in NKNT-3/NTCP #28.3.25.13 cells

Since high-level expression of STING mediated HBV-triggered type III IFN induction in NKNT-3/NTCP #28.3.25.13 cells (Figs. 4D and 4E), we next examined whether high-level expression of STING was implicated in the induction of not only type III IFN but also pro-inflammatory cytokine including IL-6 through the NF-κB signaling pathway. *IL-6* mRNA induction in p-dGdC-transfected NKNT-3/NTCP #28.3.25.13 cells was higher than that in p-dGdC-transfected NKNT-3/NTCP #28.3.8 cells (Fig. 5A). In addition, the knockdown of STING reduced *IL-6* mRNA induction in p-dGdC-transfected NKNT-3/NTCP #28.3.25.13 cells (Fig. 5B). Since the phosphorylation of NF-κB p65 at Ser536 was required for the activation of noncanonical NF-κB signaling pathway [17], we next compared the phosphorylation of NF-κB p65 at Ser536 between p-dGdC-transfected NKNT-3/NTCP #28.3.8 cells and #28.3.25.13 cells. Our results indicated that NF-κB p65 was phosphorylated at Ser536 in p-dGdC-treated NKNT-3/NTCP #28.3.25.13 cells, but not #28.3.8 cells (Fig. 5C). These results suggest that high-level expression of STING enhances p-dGdC-triggered *IL-6* mRNA induction through the noncanonical NF-κB signaling pathway in NKNT-3/NTCP #28.3.25.13 cells. We next examined whether HBV infection also triggered *IL-6* mRNA induction through the noncanonical NF-κB signaling pathway in NKNT-3/NTCP #28.3.25.13 cells. Interestingly, HBV infection, but not mock or UV-HBV infection, triggered the phosphorylation of NF-κB p65 at Ser536 (Fig. 5D) and subsequently induced *IL-6* mRNA (Fig. 5E) in NKNT-3/NTCP #28.3.25.13 cells. These results suggest that high-level expression of STING is implicated in HBV-triggered pro-inflammatory cytokine induction through the noncanonical NF-κB signaling pathway in NKNT-3/NTCP #28.3.25.13 cells. NKNT-3/NTCP #28.3.25.13 cells are a useful tool for studying hepatic carcinogenesis caused by the HBV-triggered inflammatory response through the NF-κB signaling pathway.

**Figure 5.**
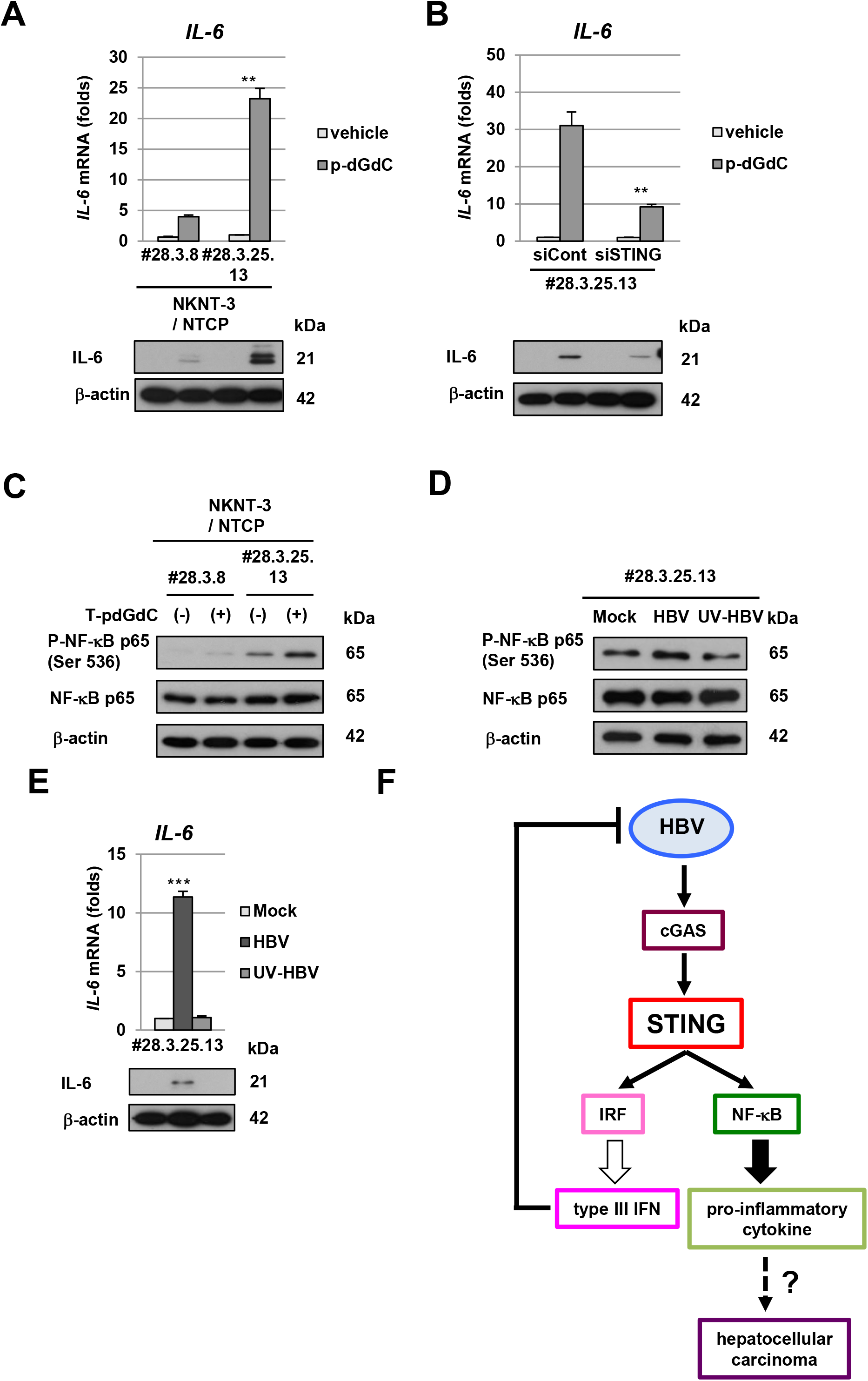
High-level expression of STING was required for HBV-triggered inflammatory response in NKNT-3/NTCP #28.3.25.13 cells. A (upper panel) Quantitative RT-PCR analysis of *IL-6* mRNA in p-dGdC-transfected NKNT-3/NTCP #28.3.8 or #28.3.25.13 cells. Each mRNA level was calculated relative to the level in vehicle-transfected NKNT-3/NTCP #28.3.25.13 cells, which was set at 1. \*\**P* < 0.01 versus p-dGdC-transfected NKNT-3/NTCP #28.3.8 cells. (lower panel) Western blot analysis of IL-6 in p-dGdC-transfected NKNT-3/NTCP #28.3.8 or #28.3.25.13 cells. The cell lysate was prepared as described in Fig. 3C. B (upper panel) Quantitative RT-PCR analysis of *IL-6* mRNA in p-dGdC-transfected NKNT-3/NTCP #28.3.25.13 siCont cells or NKNT-3/NTCP #28.3.25.13 siSTING cells. Each mRNA level was calculated relative to the level in vehicle-transfected NKNT-3/NTCP #28.3.25.13 siCont cells, which was set at 1. \*\**P* < 0.01 versus p-dGdC-transfected NKNT-3/NTCP #28.3.25.13 siCont cells. (lower panel) Western blot analysis of IL-6 in p-dGdC-transfected NKNT-3/NTCP #28.3.25.13 siCont cells or NKNT-3/NTCP #28.3.25.13 siSTING cells. The cell lysate was prepared as described in Fig. 3C. C Western blot analysis of phosphorylated NF-κB p65 at Ser536 in p-dGdC-transfected NKNT-3/NTCP #28.3.8 or #28.3.25.13 cells. D Western blot analysis of phosphorylated NF-κB p65 at Ser536 in mock-, HBV-, or UV-HBV-infected NKNT-3/NTCP #28.3.25.13 cells. E (upper panel) Quantitative RT-PCR analysis of *IL-6* mRNA in mock-, HBV-, or UV-HBV-infected NKNT-3/NTCP #28.3.25.13 cells. Each mRNA level was calculated as described in Fig. 3D. \*\*\**P* < 0.001 versus mock- or UV-HBV-infected NKNT-3/NTCP #28.3.25.13 cells, respectively. (lower panel) Western blot analysis of *IL-6* mRNA in mock-, HBV-, or UV-HBV-infected NKNT-3/NTCP #28.3.25.13 cells. F Proposed model of the HBV-triggered host innate immune response and inflammatory response through STING.

## Discussion

Cytoplasmic DNA or RNA sensors trigger the innate immune responses and the inflammatory responses by recognizing viral PAMPs. We previously reported that one of the cytoplasmic DNA sensors, cGAS, recognized HBV DNA as viral PAMPs and subsequently induced the innate immune response through its adaptor protein, STING [10]. In the present study, we found that the immortalized human hepatocyte NKNT-3 cells exhibited HBV susceptibility by stably expressing the exogenous NTCP (Figs. 1C and 1D). Cells of one of the NKNT-3/NTCP cell-derived clones, NKNT-3/NTCP #28.3.25.13, highly expressed STING and exhibited resistance to HBV through STING-mediated type III IFN induction (Figs. 4C, 4E, and 4F). Interestingly, STING was highly phosphorylated in p-dGdC-transfected NKNT-3/NTCP #28.3.25.13 cells, but not in the parent, #28, #28.3, or #28.3.8 cells (Fig. 4D). However, it is uncertain why the expression and phosphorylation levels of STING differed among the NKNT-3/NTCP cell-derived cell clones. In humans, several single nucleotide polymorphisms (SNP) of STING have been discovered [18]. SNPs of STING have been shown to cause autoinflammatory diseases such as STING-associated vasculopathy with onset in infancy [19] and familial chilblain lupus [20]. These SNPs are implicated in the dysregulation of host innate immune responses and inflammatory responses through a loss-of-function mutation or a gain-of-function mutation of STING. Further analysis is needed to identify the gain-of-function mutation(s) in STING in NKNT-3/NTCP #28.3.25.13 cells.

In the present study, we showed that HBV infection induced type III IFN, but not IFN-β (type I IFN), through a STING-mediating signaling pathway in NKNT-3/NTCP #28.3.25.13 cells (Fig. 5F). Sato et al. previously reported that a cytoplasmic RNA sensor, RIG-I, recognized HBV pgRNA and subsequently induced type III but not type I IFN through its adaptor protein, IPS-1, in human primary hepatocytes [21]. These results suggest that HBV suppresses the induction of type I IFN but not type III IFN. One of the HBV proteins, HBV polymerase, suppressed STING-mediated IFN-β induction by disrupting K63-linked ubiquitination of STING [22]. Another study also reported that HBx bound IPS-1 and suppressed the activation of IFN-β [23]. However, in these studies, it was unclear whether HBV suppressed the induction of type III IFN through these HBV proteins. Our results showed that HBV transiently induced *ISG56* mRNA induction at 5 and 9 days, but not at 13 days, after HBV infection in NKNT-3/NTCP #28.3.25.13 cells (Fig. 3D). This result suggests that HBV possesses two opposite functions to simultaneously trigger or suppress the induction of type III IFN. Further analysis is needed to examine whether or not HBV suppresses the induction of type III IFN.

We also showed that HBV infection induced a pro-inflammatory cytokine, IL-6, through the noncanonical NF-κB signaling pathway in NKNT-3/NTCP #28.3.25.13 cells (Fig. 5F). STING also mediates host inflammatory responses by triggering its downstream NF-κB signaling pathway [8, 9]. A STING-triggered host inflammatory response has been reported to be associated with hepatic diseases [24, 25]. In nonalcoholic fatty liver disease, STING promotes hepatocyte injury by inducing inflammation [24]. In addition, STING mediates liver injury and fibrosis in mice administered CCl4 (a chemical inducer of hepatocyte death) [25]. Moreover, based on the results of several previous studies, STING is also thought to play an important role in tumor development [26]. Interestingly, STING may exert two opposite effects (tumor-suppressing and tumor-promoting effects) on tumor development under different situations. For example, in breast cancer, STING and its downstream signaling may suppress the tumor or the cancer metastasis [27, 28]. In contrast, STING is also required for cell survival and regrowth in breast cancer [29, 30]. However, the results of the present study do not clarify whether the HBV-triggered NF-κB signaling pathway causes liver diseases and tumor development. Further analysis will also be needed to examine how HBV causes liver diseases and finally HCC through a STING-mediated NF-κB signaling pathway.

In the present study, we established a novel HBV infectious cell culture model by using NKNT-3 cells. Since NKNT-3 cells exhibit a non-neoplastic phenotype [12], our HBV infectious cell culture model is expected to be a useful tool for the study of hepatic carcinogenesis caused by HBV-triggered innate immune responses and inflammatory responses.

## Materials and Methods

### Cell culture

Human immortalized hepatocyte NKNT-3 cells, which were kindly provided by N. Kobayashi and M. Namba (Okayama University). Human hepatoma HepG2/NTCP cells were cultured as previously described [10]. HepG2.2.15 Cont, HepG2.2.15 cGAS/STING, and HepG2.2.15 cGAS GSAA/STING cells were maintained in medium including blasticidin and puromycin as previously described [10].

### Establishment of an NKNT-3 cell line stably expressing exogenous NTCP and the derivation of its cell clones

NKNT-3 cells stably expressing exogenous NTCP (designated NKNT-3/NTCP cells) were established as previously described [10]. NKNT-3/NTCP-derived cell clones were isolated from their parental cells by the limited dilution method. We evaluated HBV susceptibility by HBV/NLuc assay [13] and, from the several tens of cell clones obtained, selected a cell clone exhibiting susceptibility or resistance to HBV. By repeating the cell cloning and selection process, we obtained cell clones exhibiting the different levels of susceptibility to HBV

### HBV/NLuc assay

HBV/NLuc was prepared as previously reported [13]. Intracellular NLuc activity was measured at 5, 9, and 13 days after the inoculation of HBV/NLuc. For the measurement of NLuc activity, we used a Nano-Glo luciferase assay system (Promega, Madison, WI, USA). Data are the means ± SD from three independent experiments.

### Western blot analysis

Western blot analysis was performed as previously described [31]. Anti-Myc (PL14; Medical & Biological Laboratories, Nagoya, Japan), anti-ISG15 (H-150; Santa Cruz Biotechnology, Dallas, TX, USA), anti-ISG56, anti-cGAS, anti-phospho-STING (Ser366), anti-STING, anti-phospho-NF-κB p65 (Ser536), anti-NF-κB p65 (Cell Signaling Technology, Beverly, MA, USA), and anti-β-actin (AC-15; Sigma-Aldrich, St. Louis, MO, USA) were used as primary antibodies.

### Flow cytometric analysis

Cell surface expression of exogenous NTCP was detected by a flow cytometer as previously reported [32]. Anti-Myc (PL14; Medical & Biological Laboratories), and FITC-conjugated goat anti-mouse antibody (Jackson ImmunoResearch Laboratories, West Grove, PA, USA) were used as primary and secondary antibody, respectively.

### Analysis of HBV RNA

HBV was prepared from the supernatant of HepG2.2.15 cells as previously reported [10]. Cells were infected with HBV at 10^3^ HBV genome equivalents per cell, unless otherwise described. For the analysis of intracellular HBV RNA after the infection of HBV, we performed quantitative RT-PCR analysis and Northern blot analysis as previously reported [10].

### Quantitative RT-PCR analysis

At 5, 9, and 13 days after HBV inoculation or at 6 hours after the transfection of an *in* viŕrosynthesized ligand, p-dGdC (Invivogen, San Diego, CA, USA), we performed quantitative RT-PCR analysis as previously described [33]. For quantitative RT-PCR analysis, we used primer sets previously described for ISG56 [34], IFN-β [34], cGAS [10], STING [10], IL-6 [33], and GAPDH [33]. We also prepared forward and reverse primer sets for IFN-λ1 (5’-CTGGGAAGGGCTGCCACATT-3’ (forward) and 5’-TTGAGTGACTCTTCCAAGGCG-3’ (reverse)) and IFN-λ2/3 (5’-CAGCTGCAGGTGAGGGAG-3’ (forward) and 5’-CTGGGTCAGTGTCAGCGG-3’ (reverse)).

### RNA interference

The day after mock or HBV infection, we introduced small interfering RNAs (siRNAs) targeting STING or nontargeting siRNAs into NKNT-3/NTCP #28.3.25.13 cells as previously described [35]. At 4 days after the introduction of siRNAs, we isolated the total RNA or cell lysate, and subjected it to quantitative RT-PCR analysis or western blot analysis, respectively.

### Generation of cells stably expressing exogenous STING

To construct pCX4bleo/HA-STING retroviral vector, we introduced STING (accession no. NM_198282) cDNA containing a full-length ORF into the pCX4bleo/HA retroviral vector as previously reported [36]. pCX4bleo/HA-STING I200N [16] was also constructed using PCR mutagenesis with primers containing base alterations. These vectors were introduced into NKNT-3/NTCP #28.3.8 cells by retroviral transfer and then the cells stably expressing exogenous STING or STING I200N were selected by Zeocin (Thermo Fisher Scientific, Carlsbad, CA, USA).

### Statistical analysis

Statistical analysis was performed to determine the significance of differences among groups by using Student’s ŕ-test. *P* < 0.05 was considered statistically significant.

## Acknowledgements

We thank Marie Iwado, Masayo Takemoto, and Masato Ono for their technical assistance. We also thank Dr. Tsuyoshi Akagi for the provision of pCX4bleo retroviral vectors. This research was supported by the Japan Agency for Medical Research and Development (AMED) under grant numbers JP17fk0310107 and JP17fk0310103, and by a grant from the Wesco Foundation.

## Author Contributions

HD and NK designed the research. HD performed most of the experiments. HI contributed pCX4bleo HA-STING I200N. NK performed the cell cloning by the limited dilution method. HD, HI, YU, SS and NK analyzed the data. HD wrote the paper. All authors reviewed the manuscript.

